# Remodelling of pSK1 Family Plasmids and Enhanced Chlorhexidine Tolerance in Methicillin-Resistant *Staphylococcus aureus*

**DOI:** 10.1101/457838

**Authors:** Sarah L Baines, Slade O Jensen, Neville Firth, Anders Gonçalves da Silva, Torsten Seemann, Glen Carter, Deborah A. Williamson, Benjamin P Howden, Timothy P Stinear

## Abstract

*Staphylococcus aureus* is a significant human pathogen whose evolution and adaptation has been shaped in part by mobile genetic elements (MGEs), facilitating global spread of extensive antimicrobial resistance. However, our understanding of the evolutionary dynamics surrounding MGEs is incomplete, in particular how changes in the structure of multidrug-resistant (MDR) plasmids may influence important staphylococcal phenotypes. Here, we undertook a population-and functional-genomics study of 212 methicillin-resistant *S. aureus* (MRSA) ST239 isolates collected over 32 years to explore the evolution of the pSK1 family of MDR plasmids, illustrating how these plasmids have co-evolved with and contributed to the successful adaptation of this persistent MRSA lineage. Using complete genomes and temporal phylogenomics we reconstructed the evolution of the pSK1 family lineage from its emergence in the late 1970s, with multiple structural variants arising. Plasmid maintenance and stability was linked to IS*256*- and IS*257*-mediated chromosomal integration and disruption of plasmid replication machinery. Overlaying genomic comparisons with phenotypic susceptibility data for gentamicin and chlorhexidine, it appeared that pSK1 has contributed to enhanced resistance in ST239 MRSA through two mechanisms: (i) acquisition of plasmid-borne resistance mechanisms increasing rates of gentamicin resistance and reduced chlorhexidine susceptibility, and (ii) changes in plasmid configuration linked with further enhancement of chlorhexidine tolerance. While the exact mechanism of enhanced tolerance remains elusive, this research has uncovered a clear evolutionary response of ST239 MRSA to chlorhexidine, one which may contribute to the ongoing persistence and adaptation of this lineage within healthcare institutions.

## Importance

The biocide chlorhexidine is fundamental to infection control practices which prevent nosocomial infection and it is highly effective for decolonisation of *S. aureus* from human skin. There have been increasing reports of staphylococcal populations evolving tolerance to chlorhexidine, suggesting that the increasing use and reliance on this biocide may provide a significant selection pressure influencing the evolution of staphylococcal populations. Chlorhexidine tolerance in *S. aureus* is commonly enabled by the acquisition of an efflux pump, however alternative mechanisms influencing tolerance are poorly understood. In this study we demonstrate a previously unrecognised phenomenon, plasmid structural remodelling, by which *S. aureus* may developed enhanced chlorhexidine tolerance. Further, we highlight the importance of undertaking a detailed exploration of the evolutionary dynamics surrounding mobile genetic elements, which contribute immensely to the evolution of microbial species and provide here a framework for how such an analysis can be performed.

## Introduction

Mobile genetic elements (MGEs) play a central role in microbial evolution; serving as a mechanism by which genetic material can be transferred, disseminated and rearranged, allowing for rapid adaptation to new and changing environments. Nowhere is this more apparent than in the global dissemination of genes encoding mechanisms of antimicrobial resistance and virulence in populations of clinically significant bacteria [1-4]. *Staphylococcus aureus* is a leading cause of bacterial infections in humans and invasive staphylococcal disease is associated with significant morbidity and mortality [5, 6]. One of the oldest and truly pandemic lineage of *S. aureus* is multilocus sequence type (ST) 239; a multidrug-resistant, healthcare associated (HA) methicillin-resistant *S. aureus* (MRSA) clone first identified in the late 1970s [7-9]. Multiple studies have used genomics to explore the evolution of ST239 MRSA, revealing mechanisms which have contributed to its global spread, extensive antimicrobial resistance repertoire, and persistence in healthcare environments [10-15]. In Australia, ST239 has been the dominant HA-MRSA lineage for nearly four decades (national surveillance reports:http://agargroup.org.au/agar-surveys/). Although its prevalence is declining in the regions, having recently been surpassed by the epidemic EMRSA-15 (ST22) clone [16], ST239 is still regularly recovered as a cause of invasive disease in multiple Australian states [17]. We have previously described the long-term evolution of ST239 MRSA in Eastern-Australian hospitals, one of convergent and adaptive evolution of two genetically distinct ST239 clades towards increased antimicrobial resistance at the cost of attenuated virulence [15]. This initial work largely focused on changes occurring within conserved regions of the genome with limited exploration of the accessory genome that is primarily composed of MGEs.

Early ST239 MRSA were first recognised in Australia because they displayed resistance to gentamicin [18-20], encoded for by a bifunctional acetyltransferase-phosphotransferase gene *aac(6’)-aph(2”)* commonly carried on pSK1-like plasmids as part of the composite transposon (Tn)*4001* [21, 22]. The focus of multiple studies, pSK1 represents a family of intermediate sized (20-40 kb), theta-replicating staphylococcal plasmids that have been recovered in Australia and the United Kingdom [21, 23-26]. Multiple antibiotic resistance mechanisms are variably encoded for on pSK1-like plasmids. In addition to *aac(6’)-aph(2”),* resistance to penicillinase-labile penicillins is encoded for by *blaZ* as part of Tn*552* [24, 25], and trimethoprim resistance is mediated by an insensitive dihydrofolate reductase (dfr), encoded by *dfrA* and carried as part of *Tn4003* which represents a cointegrated remnant of a pSK639-like plasmid [27, 28]. Additionally, harboured by pSK1-like plasmids is a *qacA* gene encoding a quaternary ammonium compound (QAC) multidrug-efflux pump [29], which can mediate tolerance to cationic biocides, most notably chlorhexidine [30]. The presence of a plasmid-borne *qacA* gene, and predicted biocide tolerance, has previously been attributed to an outbreak of ST239 MRSA in the United Kingdom in the early 2000s [31, 32].

The divalent cationic biocide chlorhexidine digluconate (CHX) was first described in 1954 and is a fundamental component of infection control practices to prevent nosocomial infections [33, 34]. It is one of the most widely used antiseptic agents because of it broad spectrum of activity against bacteria, fungi, and enveloped viruses, as well as its good safety record and general tolerability [35-37]. Resistance at in-use concentrations, typically a 0.2 to 4.0% CHX solution in water, has not been reported in *S. aureus*, however the phenomena of enhanced tolerance appears to be increasingly common [38, 39], and is reviewed in [40]. CHX tolerance, defined here as an increase in the minimum inhibitory concentration (MIC) and/or minimum bactericidal concentration (MBC) to a level that remains below in-use concentrations, has been associated with the acquisition or mutation of cationic biocide active efflux pumps, the most commonly reported being the QAC efflux systems [30, 41, 42]. Most reports of staphylococcal populations evolving CHX tolerance demonstrated either a phenotypic shift in MIC/MBC or detected an increase in the prevalence of genes encoding efflux systems [40, 43]. In *S. aureus* this is mainly associated with *qacA,* often referred to as to *qacA/B* as the encoded peptides differs by only a single amino acid [30, 40, 44]. It is important to note that acquisition of a *qac* gene does not consistently lead to phenotypic tolerance, nor is an increase in CHX tolerance invariably associated with an efflux system [40, 45, 46]. Subsequently, shifts in the prevalence of *qac* genes in staphylococcal populations may not correspond with the development of CHX tolerance. While several studies do combine phenotypic and molecular data, this work has largely been performed on clonally-diverse or genomically-undefined populations, and thus the evolutionary mechanisms responsible are often unclear [38, 43].

Here we present a detailed exploration of the evolutionary dynamics surrounding the pSK1 family of plasmids, focusing on a single well-defined staphylococcal lineage, the ST239 MRSA population circulating in Australia. The primary aims of this work were to: (i) identify and understand the mechanisms of adaptation with a specific focus on changes in the structural configuration of pSK1-like plasmids, and (ii) explore the phenotypic consequences of these changes in ST239 MRSA, with particular interest in the development of enhanced CHX tolerance.

## Results and Discussion

In Australia, the ST239 MRSA population is composed of two genetically distinct clades [15]. The oldest clade is largely restricted to Australia and represents the original lineage circulating in the region. Conversely, the newer clade represents an intercontinental transmission event, introducing the lineage circulating in South East Asia into Australia, a more recent estimate placing this event in the late 1990s (Figure S1). These ST239 clades were originally referred to as Clade 1 and Clade 2 [15], however in this publication we have renamed them as the Australian clade (original) and the Asian-Australian clade (introduced), respectively.

Plasmids related to pSK1 were first identified in ST239 MRSA circulating in Australia in the early 1980s [21, 23, 24]. An *in silico* examination of contemporary isolates revealed that pSK1-like plasmids are still maintained in this population (Figure S1), suggesting the extended co-evolution of pSK1-like plasmids and ST239 MRSA and hence prolonged exposure to the highly selective healthcare environment. To explore the evolutionary dynamics surrounding the pSK1 family we undertook a detailed phenotypic and comparative genomic analysis of 212 temporally (1980 to 2012) and geographically diverse Australian ST239 MRSA. To aid in understanding the changes in plasmid configuration which are described in this publication the structure of the family prototype, pSK1, is illustrated in Figure 1A and described in the Supplementary Results.

**Figure 1.**
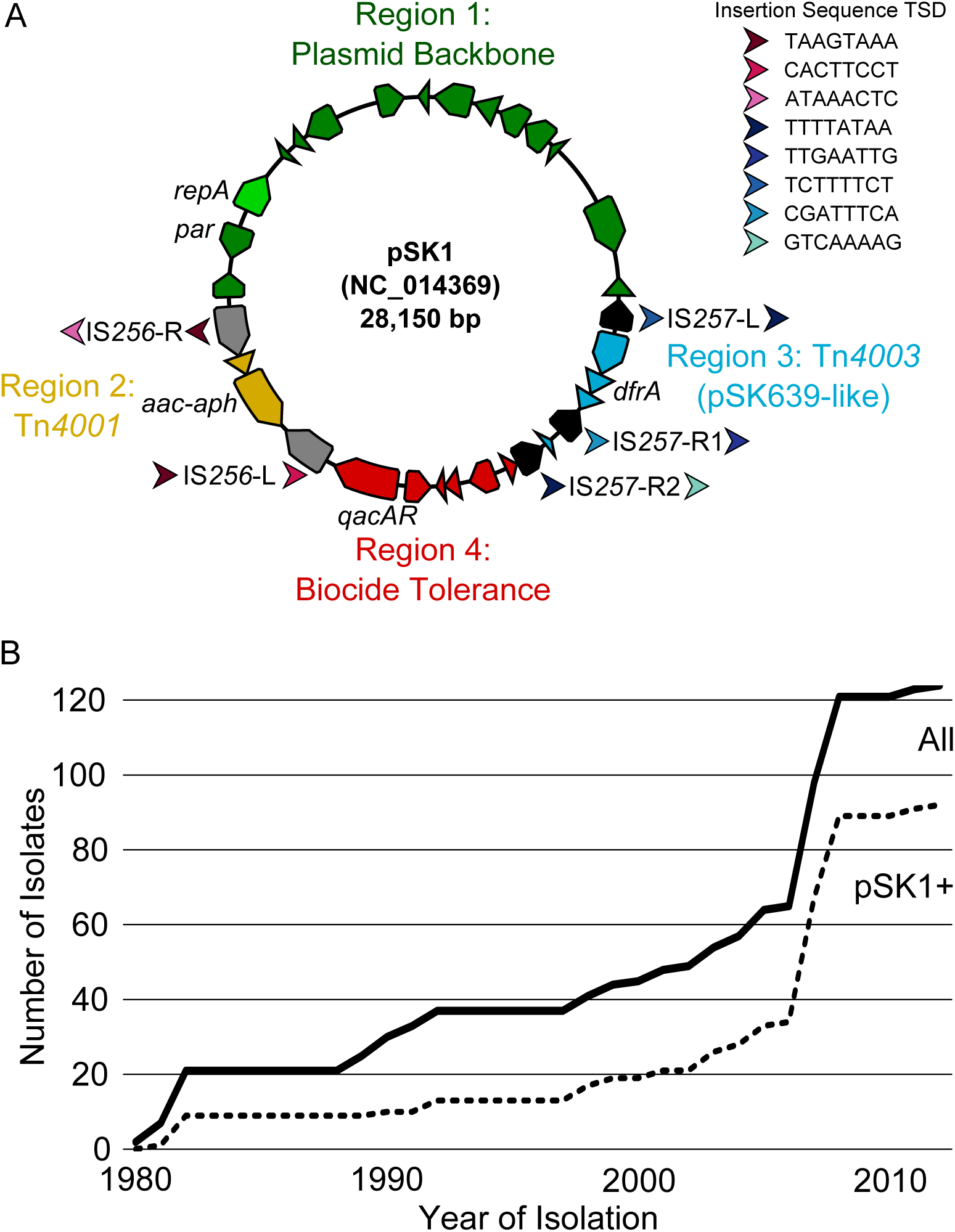
Structure and prevalence of pSK1 plasmid s in ST239 MRSA. (A) pSK1 sequence and annotations are that previously published, GenBank accession GU565967.1 [47]. Predicted CDS have been coloured based on defined regions. Insertion sequences (IS) are coloured grey (IS*256*) and black (IS*257*), with target site duplications (TSD) illustrated: arrows indicate upstream/downstream sequences, orientation, and are coloured to represent unique sequences (refer to key). (B) Graph illustrates the increasing prevalence of pSK1 family plasmids in the Australian ST239 clade overtime. The cumulative total of the sampled population is indicated by the solid line and the proportion in which a pSK1-like ike plasmid was identified by the broken line.

### Prevalence and Structural Variability of pSK1 Family Plasmids

To identify pSK1-like plasmids, the presence and synteny of plasmid genes was assessed *in silico.* This analysis found that pSK1-like plasmids were solely carried by isolates belonging to the Australian clade, present in 92/124 isolates (74.1%). A comparison of plasmid carriage with year of isolation illustrated that the prevalence of pSK1-like plasmids had increased over time (Figure 1B), suggesting an evolutionary benefit for their acquisition and maintenance in this population. There was no evidence of plasmid sharing between the Australian and Asian-Australian clades (Figure 2). However, this finding was not surprising due to the geographic separation of the two clades; the Australian clade has been predominantly recovered in the states of New South Wales and Queensland since the early 2000s, and the newer Asian-Australian clade has almost exclusively been recovered in Victoria since its introduction [15] (Figure 2A).

**Figure 2.**
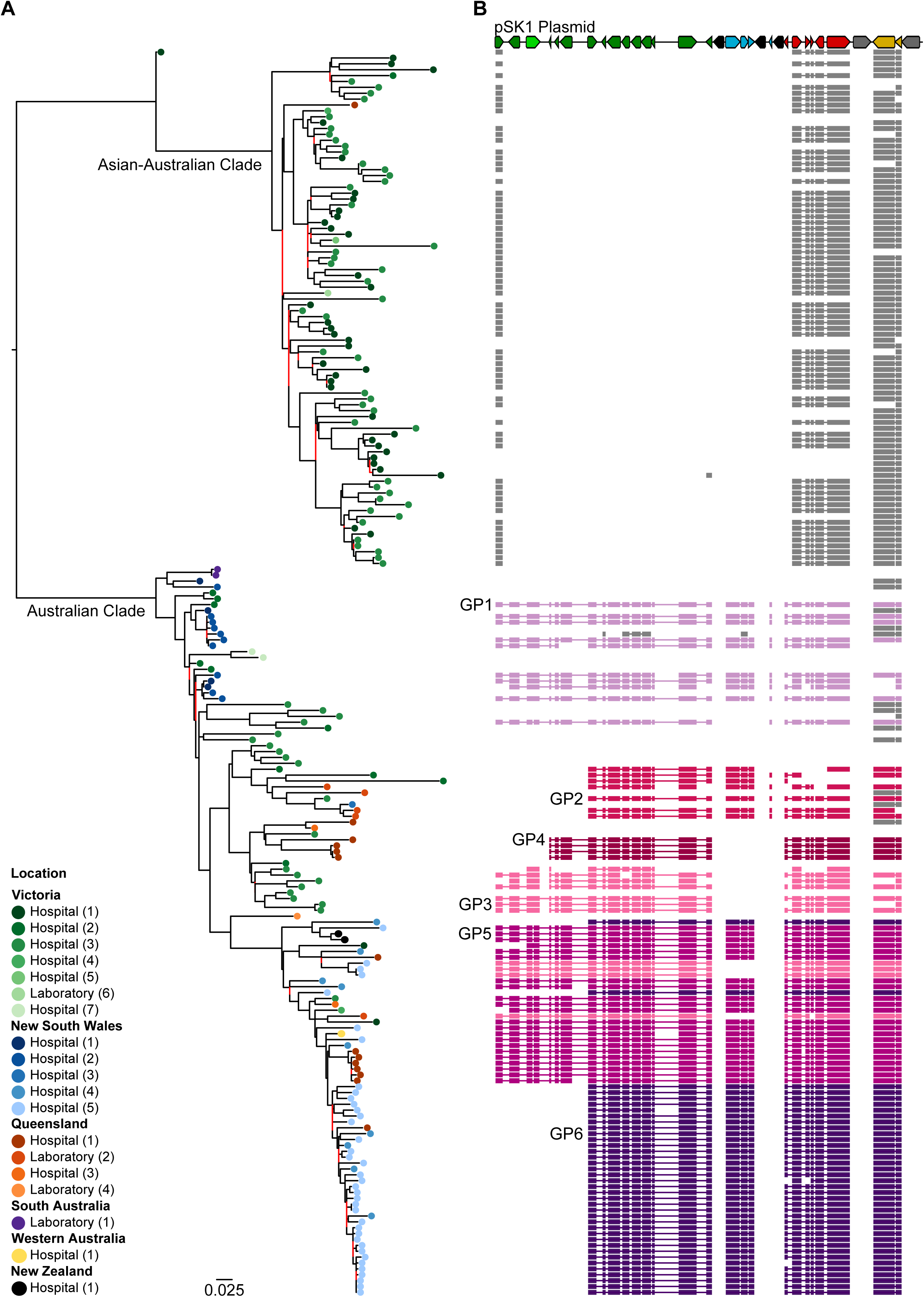
pSK1 plas mid gene presence and synteny. (A) Maximum likelihood phylogenetic tree inferred from 3,883 core genome SNPs illustrates the population structure of ST239 *S. aureus* in Australia. Tips are coloured based on location (refer to key). Branches with < 70% bootstrap support are coloured red. (B) Coloured blocks represent the identification of a pSK1 gene, using a 95% amino acid homology threshold (excluding IS elements). Box length is reflective of gene length and ordered based on pSK1 (Figure 1A). Boxes are linked if CD S were found to be syntenic. Coloured boxes reflect one of six defined gene patterns (GP), with location of GP label indicating the isolates selected for long-read sequencing.

A closer examination of the 92 isolates in which a pSK1-like plasmid was identified showed considerable variation in the presence of plasmid genes (Figure 2B). When aligned to a model for the ST239 population structure it became apparent that the variation observed was phylogenetically correlated and suggested the emergence or acquisition of a limited number of pSK1-like structural variants (SVs), with subsequent clonal expansion. Six broad gene patterns (GP) could be resolved amongst these data (Figure 2B, Supplementary Results). Differences between the GPs correlated with the variable presence of the composite transposons in pSK1. For example, the absence of Tn*4003* in GP3 and GP4 (Figure 2B). Additionally, the loss of three to six syntenic genes in the plasmid backbone, including the replication initiation protein (*repA*) and the plasmid partitioning (*par*) genes (observed in GP2, GP4 and GP6, Figure 2B). This finding suggested that some of the pSK1-like SVs may be chromosomally integrated. Examination of the pSK1 gene presence and synteny in the Asian-Australian clade revealed that 5/6 syntenic genes associated with pSK1 region 4 (including *qacAR*) were highly prevalent (Figure 2B). These isolates were found to carry an alternative *qacA*-containing plasmid, specifically a pTW20_1-like plasmid [31]. Unlike pSK1, the pTW20_1-like gene structure appeared to be largely conserved. A description of this second plasmid population is provided in the Supplementary Results.

### Structural Variants of pSK1

To explore the extent of structural remodelling (defined here as changes in plasmid gene content and/or configuration) that has occurred in the pSK1-like plasmid population, long-read sequencing was conducted to establish a reference sequence for the six expected SVs. Additionally, the genome assemblies of all isolates were mined for unique plasmids features, used to determine the prevalence of each SV in the wider ST239 population. This work has been summarised below. For a more detailed explanation refer to the Supplementary Results.

Using this approach eight distinct pSK1-like plasmids were identified, having arisen largely through the activities of *IS256* and/or IS257. These changes can be classified into three categories. The first was IS-mediated gain or loss of the composite transposons Tn*4001* and Tn*4003*. This was observed in three SVs, termed SV1, SV3 and SV4 (representing GP1, GP3, and GP4, respectively). SV1 was found to lack the aminoglycoside resistance conferring Tn*4001* and is structurally equivalent to the previously identified pSK7 (Figure 3A) [24, 47]. With only a single copy of the IS*256* target site duplications (TSD) that flank Tn*4001* in pSK1 identified, it was unclear whether SV1 had lost Tn*4001* or had never acquired it. Given when the isolates in which SV1 were recovered (1980 and 1982) and their location close to the tree root in a time-aware phylogenetic model for the Australian clade (Figure S2), SV1 likely represented a progenitor of pSK1 prior to gaining Tn*4001*. Both SV3 and SV4 were found to lack Tn*4003* (Figure 3A,3B). Instead they contained only a single copy of IS*257* with the same TSD that flanked this region in pSK1, in a configuration equivalent to that of pSK14 [24, 47]. Unlike SV1, the phylogenetic location of isolates carrying SV3 and SV4 strongly supported deletion of Tn*4003* (Figure S2), potentially through homologous recombination between the flanking IS*257*s rather than historic absence. This notion is consistent with the proposed evolution of Tn*4003*, which through IS*257*-mediated transposition from a pSK639-like plasmid was acquired into a pSK1-like precursor to generate the plasmid cointegrate [28].

**Figure 3.**
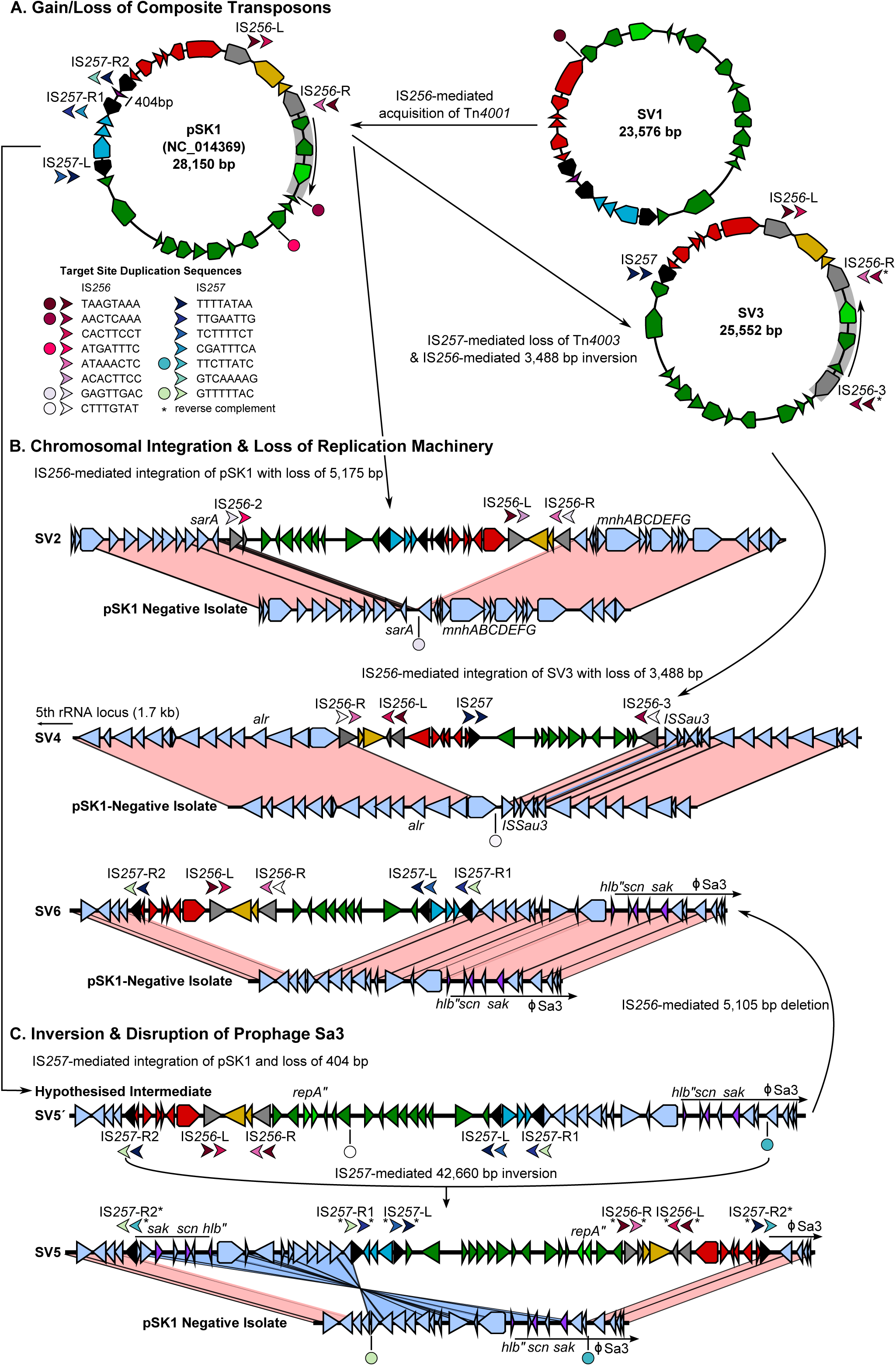
pSK1-like structural variants. Illustrated is a schematic for th e development of the pSK1-like plasmid variants. Predicted CDS are coloured based on defined pSK1 regions (Figure 1A), IS*256* are grey, IS*257* are black, and chromosomal CDS are lilac. Arrows denote upstream/downstream target site duplications (TSD) and direction denotes IS orientation. Circles represent a single copy of a TSD not adjacent to an IS element. Arrows and circles are coloured to reflect unique sequences (refer to key) and a star has been used to indicate a TSD present in the reverse complement to what was expected. In the structural comparisons, connected regions share e 98% nucleotide sequence identity, coloured pink or blue to indicate matching or reverse orientation, respectively. (A) Illustrates the emergence of pSK1-like variants through IS-mediated loss/gain of the composite transposons: (i) Tn*4001* acquired in SV1 to produce pSK1, and (ii) Tn*4003* deleted from pSK1 to produce SV3 (with an additional IS*256*-mediated inversion). (B) Illustrates three IS-mediated chromosomal integration events: (i) integration of pSK1 adjacent to *sarA* (SV2), integration of SV3 near air (SV4), and integration of pSK1 near φSa3 (SV6). All three SVs have undergone IS*256*-mediated exclusion/deletion of CDS encoding the plasmid replication mach inery. (C) Illustrates the large IS*257*-mediated inversion event in the hypothesised SV5’ th at gave rise to SV5 and resulted in the fragmentation of φSa3.

The second category was IS-mediated chromosomal integration and disruption of the plasmid replication machinery, through IS- and non-IS-mediated deletions/exclusion events. Chromosomal integration was observed in five pSK1-like SVs. Genomic island SV2 (representing GP2) emerged from IS*256*-mediated integration of pSK1 adjacent to the staphylococcal accessory regulator A (*sarA*) gene (Figure 3B). Likewise, SV4 (representing GP4) emerged from IS*256*-mediated integration of SV3 adjacent to a predicted aerobactin biosynthesis gene and in close proximity to an alanine racemase (*alr*) gene (Figure 3B). In both cases, integration was likely preceded by an IS*256* transposition event, resulting in the addition of an extra IS*256* within each plasmid precursor. Interaction between the novel and native IS*256* copies led to the formation of circular intermediates, with each composite Tn encompassing the majority of the plasmid sequence but excluding a segment of the plasmid backbone, resulting in the loss of six and three syntenic CDS in SV2 and SV4, respectively. The three remaining SVs had all arisen from a single IS*257*-mediated chromosomal integration event. Only SV5 and SV6 were represented in the complete genomes; a hypothesised structure has been proposed for the other SV, termed SV5’ as it represented the progenitor for both SV5 and SV6 (Figure 3B & 3C). In this case, chromosomal integration of pSK1 had occurred 9.2 kb upstream of a disrupted β-haemolysin gene (disruption resulting from integration of prophage Sa3) and likely involved IS*257*-mediated replicative transposition (for integration) and homologous recombination to account for the partial deletion of Tn*4003*. Genomic island SV5 had an additional 45 bp deletion in *repA* (Figure 3C), and SV6 had a deletion of six syntenic CDS from the plasmid backbone likely resulting from IS*256*-mediated homologous recombination (Figure 3B). All deletion events invariably resulted in the loss of genes, or predicted loss of function of the plasmid replication machinery, which has a known role in stabilising newly integrated elements by removing any interference with chromosomal replication [48].

The third category represented two IS-mediated inversion events. The first was an IS*257*-mediated inversion of SV5’, giving rise to genomic island SV5 (Figure 3C). This appears to be the result of intramolecular replicative transposition in the inverse orientation and resulted in the inversion of a 42.7 kb region [4]. This has reversed the orientation of all SV5 CDS and split φSa3 (Figure 3C). A second smaller IS*256*-mediated inversion event was identified in SV3. Transposition of IS*256* followed by homologous recombination between the new and a native IS*256* copy (in inverted orientation) has led to a 3.5 kb inversion in the plasmid backbone and reversed the orientation of *repA*, *par* and a hypothetical CDS (Figure 3A). Further, this is the same region deleted in SV4. While the exact consequences of these inversions are unknown the division and partial inversion of φSa3 should prevent excision of the prophage.

### Convergent Evolution of pSK1-like Variants

Analysis of unique plasmid features in the short-read *de novo* assemblies enabled assignment of all isolates harbouring a pSK1-like plasmid to one of the eight SVs and resolved all inconsistencies between the phylogenetic population model and the originally identified GPs (Supplementary Results). This finding strongly suggested that once a novel pSK1-like plasmid emerged it was maintained and structurally remodelled during clonal expansion, but rarely if ever horizontally transferred between ST239 isolates. To explore potential evolutionary patterns in the plasmid population, the SVs were overlayed on to a Bayesian-inferred time-aware phylogenetic model for the Australian clade (Figure S2).

The first pSK1-like plasmids to have appeared in the Australian ST239 clade were SV1 and pSK1, in the late 1970s, consistent with the first identified gentamicin-resistant MRSA in Australia [18-20]. From this model it was estimated that pSK1 emerged in 1979 (95% highest posterior density interval [HPDI]: 1978 – 1980), following the likely acquisition of Tn*4001* into the ancestral SV1 plasmid. In the following three decades there was the emergence of SV3 around 1991 (HPDI: 1989 – 1994), in which Tn*4003* had been deleted and the replication machinery inverted. There had been at least three independent chromosomal integration events, resulting in the emergence of SV2 around 1985 (HPDI: 1983 – 1986), SV5’ around 1994 (HPDI: 1991 – 1996), and SV4 around 2000 (HPDI: 1996 – 2004). These events likely involved pSK1 as the ancestral plasmid, or SV3 as the immediate ancestor of SV4. Inversion of SV5’ in 1999 (HPDI: 1997 – 2002) led to the emergence of SV5. In each integrated variant, the replication machinery had been disrupted through IS*256*-associated gene loss or an internal *repA* deletion. It was originally reported that region 1 was conserved in all pSK1 family plasmids [47], and for the novel extra-chromosomal SVs reported in this study this remains true. However, in the chromosomally integrated SVs only a ~9.3 kb segment of the region is conserved, demonstrating > 95% nucleotide sequence identity with *S. warneri* plasmid pPI-1 (accession AB125341.3) [47, 49]. The genes encompassed within this region encode protein products predicted to be cell-envelope associated, involved in membrane transport and potentially iron acquisition, with one gene encoding a putative Fst-like toxin as part of a Type I toxin-antitoxin system [47]. The near ubiquitous conservation of this region in the pSK1-like plasmid population suggests that these encoded products beneficial and possibly contributing to plasmid maintenance [47].

In addition to the clear stepwise evolution of the pSK1-like population, there also appeared to be evidence of convergent evolution. The three integration events have all occurred independently. Examination of the *de novo* assemblies for all 92 isolates harbouring a pSK1-like plasmid identified a further two independent deletion events amongst the SV5′ and SV5 clade (Figure S2). These findings strongly suggested that the emergence of these phylogenetically distinct but structurally similar SVs is the result of a significant but unknown evolutionary pressure acting on this pSK1-like population. There is a clear role for chromosomal integration in improving the maintenance of plasmid genes in a population, and the subsequent deletion or disruption of the plasmid replication machinery is needed for stability of the integrant [48]. In the Australian clade, almost all isolates (75/77) that have descended from an ancestral genome with an integrated pSK1-like structure have maintained it (Figure S2). It is plausible that the introduction and/or increased use of antimicrobial agents and disinfectants, specifically those for which mechanisms of resistance or tolerance are encoded for by genes harboured on pSK1, could also be contributing to the evolutionary pressure promoting plasmid maintenance and driving chromosomal integration.

### Evolving Antimicrobial Resistance and Disinfectant Tolerance

To explore the impact of pSK1 family evolution on antimicrobial resistance and disinfectant tolerance, 211 isolates underwent susceptibility testing against gentamicin, trimethoprim and chlorhexidine. Isolates representing both the Australian and Asia-Australian clades were tested to enable comparisons between the two populations and discern which trends may be associated with plasmid evolution.

#### Trimethoprim

Phenotypic resistance to trimethoprim was detected in all 211 ST239 isolates, with a MIC of > 32 mg/L (Table 1). Genotypically, 75/l23 isolates (6l.0%) of the Australian clade and all 88 isolates of the Asian-Australian clade were found to harbour a mutated or acquired *dfr* gene, the *dfrA* harboured by pSK1-like plasmids or *dfrG* respectively (Table 1). A likely mechanism of resistance could not be identified in the remaining 48 isolates from the Australian clade. The lack of phenotypic variation indicated that structural variation in the plasmid population, involving the deletion or alteration of Tn*4003*, had not impacted on trimethoprim resistance in ST239 due to an unidentified genotypic redundancy. Subsequently, trimethoprim is unlikely to be acting as a significant driver of evolution in this population.

**Table 1.** Population Distributions of Antimicrobial and Biocide Resistance Genes and Phenotypic Resistance Profiles.

|  | All ST239<br>(n = 211) | Asian-Australian<br>Clade (n = 88) | Australian Clade<br>(n = 123) | SK1 Plasmid<br>(n = 91) |
| --- | --- | --- | --- | --- |
| *Phenotypic antimicrobial resistance - median (range) mg/L* |  |  |  |  |
| **Trimethoprim (MIC)<sup>a</sup>** | > 32 (-) | - | - | - |
| **Gentamicin (MIC)<sup>a</sup>** | 32 (0.023 – 256) | 256 (0.38 – 256) | 16 (0.023 – 256) | 24 (0.032 – 256) |
| **Chlorhexidine (MIC)<sup>b</sup>** | 3 (1 – 6) | 3 (1.5 – 4) | 4 (1 – 6) | 4 (1 – 6) |
| **Chlorhexidine (MBC)<sup>b</sup>** | 6 (2 – 16) | 6 (3 – 12) | 8 (2 – 16) | 8 (2 – 16) |
| *Prevalence of acquired resistance genes<sup>c</sup> (n)* |  |  |  |  |
| **dfr genes** | *dfrA* (76) | - | *dfrA* (76) | *dfrA* (76) |
|  | *dfrG* (88) | *dfrG* (88) | - | - |
| **AME genes** | *aac6'-aph2''* (180) | *aac6'-aph2''* (82) | *aac6'-aph2''* (98) | *aac6'-aph2''* (85) |
|  | *aadD* (72) | - | *aadD* (72) | *aadD* (67) |
|  | *aph(3')-III* (95) | *aph(3')-III* (87) | *aph3'-III* (8) | *aph3'-III* (5) |
| **qac genes** | *qacA* (156) | *qacA* (67) | *qacA* (89) | *qacA* (89) |
|  | *qacC* (2) | - | *qacC* (2) | - |

Abbreviations: MIC, minimum inhibitory concentration; MBC, minimum bactericidal concentration
<sup>a</sup> Phenotypic susceptibility against trimethoprim and gentamicin was determined by E-test.
<sup>b</sup> Phenotypic susceptibility against chlorhexidine was determined by broth microdilution.
<sup>c</sup> Identification of resistance genes was determined by local alignment, minimum alignment of 70% gene length and > 95% nucleotide homology required to call a match.

#### Gentamicin

Phenotypic resistance to gentamicin was detected in 185 (87.7%) isolates (Table 1). Three acquired genes encoding aminoglycoside modifying enzymes (AME) were identified [50]: (1) the bifunctional *aac(6’)-aph(2”),* found in Tn*4001* and other chromosomal and phage locations [51-53]; (2) an adenyltransferase gene *(aadD,* also known as ANT(4’)-Ia) [54, 55]; (3) a phosphotransferase gene *(aph(3’)-IIIa)* [56], which has been found co-located with *aac(6’)-aph(2”)* in φSPβ-like in ST239 *S. aureus* TW20 [31], an isolate closely related to the Asian-Australian clade [15]. The bifunctional AME was common amongst both the Australian and Asian-Australian clades, identified in 98 (79.7%) and 82 (93.2%) isolates, respectively. Conversely, the distribution genes encoding the monofunctional AMEs were more distinct (Table 1). Phenotypically the Asian-Australian clade demonstrated a significant higher average MIC to gentamicin compared to the Australian clade (*p* < 0.0001, Table S1). However, as most isolates harboured multiple AMEs, phenotypic resistance could not be attributed to a single gene. That being said, the distribution of these genes suggested that *aph(3’)-IIIa* is likely responsible for the high level resistance observed in the Asian-Australian clade, with *aac(6’)-aph(2”)* and *aadD* contributing only low level resistance in the Australian clade (Table S1).

To assess the impact of pSK1 evolution on gentamicin MIC, the phenotypic data was investigated for temporal trends. These analyses suggested a trend of increasing gentamicin MIC overtime. However, this was only observed in the Australian clade and not the Asian-Australian clade when modelled separately (Figure S3). This finding is consistent with what has been previously observed in ST239 MRSA with the glycopeptides and daptomycin, hypothesised to be the result of two evolutionary phenomena: (i) the introduction of the more resistant Asian-Australian clade into the region with successful local expansion and (ii) adaptive evolution within the Australian clade, collectively shifting the phenotype of the population overtime [15]. These same phenomena have likely contributed to the shift in gentamicin MIC. The Asian-Australian clade had already developed high level gentamicin resistance prior to arriving in Australia, through the acquisition of φSPβ-like carrying both *aac(6’)-aph(2”)* and *aph(3’)-IIIa.* Concurrently, the Australian clade had undergone adaptive evolution, with the acquisition of *aac(6’)-aph(2”)* and/or *aadD* contributing to a significant increase in MIC (Table S1); the uptake of pSK1 serving as one mechanism by which *aac(6’)-aph(2”)* could be acquired. However, plasmid evolution, with the emergence of the pSK1-like variants, appeared to have no effect on gentamicin MIC, with no temporal trend having been detected amongst the pSK1-like plasmid containing population (Figure S3, Table S2). Therefore, gentamicin also does not appear to be acting as a significant driver of plasmid structural remodelling.

#### Chlorhexidine

Phenotypic tolerance to CHX was detected in 150 (71.1%) isolates, defined as an MIC > 2 mg/L [40]. The MIC values of the population ranged from 1 - 6 mg/L, and the MBC values from 2 - 16 mg/L (Table 1). A total of 156 (73.9%) isolates were found to harbour a *qacA* gene and two a *qacC* gene (Table 1). The latter, also known as *qacD, smr* or *ebr*, although encoding a biocide active efflux pump is not active against CHX [40]). The *qacA* gene was consistently associated with isolates that carried either a pSK1-like (Australian clade) or pTW20_1-like plasmid (Asian-Australian clade). The presence of *qacA* was significantly associated with an elevated MIC and MBC to CHX (*p* < 0.0001, Table 2). Although *qacA* was equally prevalent in both the Australian and Asian-Australian clades, 89 (72.4%) and 67 (76.1%) isolates respectively, the former population demonstrated a significantly higher average MIC (*p* < 0.0001, Table 2). This phenotypic disparity remained when isolates that did not carry either plasmid were excluded from the comparison, suggesting that genotypic differences between the two plasmid populations could be contributing to the observed phenotypic variation. A comparison of all *qacAR* sequences to reference JKD6008 (Australian clade) identified six SNPs resulting in missense mutations, five in *qacA* and one in *qacR.* Examination of the prevalence, phylogenetic correlation, and phenotypic association of the mutations found that none could explain the difference in tolerance observed between the two clades (Supplementary Results). Using sequence read coverage data, plasmid copy number was also investigated. This analysis revealed that both pSK1-like and pTW20_1-like plasmids were low copy number (average of one and three copies, respectively) and maintained only a single copy of *qacA* per plasmid (Supplementary Results). While this finding was consistent with previous reports and pSK1-like variants commonly being chromosomally integrated [15], it did not explain the observed phenotypic variation and suggested that genotypic difference occurring outside of *qacAR* were responsible.

**Table 2.** Investigation of Chlorhexidine Tolerance in Australian ST239 MRSA

| Isolate Populations (n) | Median (mg/L) |  | Isolate Populations (n) | Median (mg/L) |  | P value |
| --- | --- | --- | --- | --- | --- | --- |
|  | MIC; MBC |  |  | MIC; MBC |  | MIC; MBC |
| All ST239 (211) |  |  |  |  |  |  |
| qacA Negative (55) | 0.90; 2.30 | vs | qacA Positive (156) | 3.70; 8.00 |  | < 0.0001; < 0.0001 |
| Australian Clade (123) | 3.40; 7.30 | vs | Asian-Australian Clade (88) | 2.30; 6.70 |  | < 0.0001; NS |
| SK1 plasmid (91) | 4.20; 8.60 | vs | pTW20_1-like (67) | 3.00; 7.20 |  | < 0.0001; 0.0021 |
| Asian-Australian Clade (88) |  |  |  |  |  |  |
| qacA Negative (21) | 1.90; 5.20 | vs | qacA Positive (67) | 3.00; 7.20 |  | < 0.0001; < 0.0001 |
| Australian Clade (123) |  |  |  |  |  |  |
| qacA Negative (34) | 1.80; 4.60 | vs | qacA Positive (89) | 4.30; 8.60 |  | < 0.0001; < 0.0001 |
| Extra-chromosomal SVs (17) | 3.20; 7.80 | vs | Integrated SV (74) | 4.50; 8.74 |  | 0.0086; NS |
| Integrated SV + repA deletion (25)<sup>b</sup> | 3.70; 8.50 | vs | Integrated SV + R1 deletion (49)<sup>c</sup> | 4.70; 8.60 |  | NS; NS |

Abbreviations: MIC, minimum inhibitory concentration; MBC, minimum bactericidal concentration; R1, pSK1 region 1; SV, structural variant.
<sup>a</sup> Population includes SV5' and SV5; <sup>b</sup> Population includes SV2, SV4, and SV6.

As with gentamicin, linear models were developed to investigate the CHX phenotypic data for temporal trends, which may indicate a role for pSK1 family evolution in the development of CHX tolerance (Figure S3). These models also suggested a trend of increasing MIC and MBC to CHX overtime. Again, this trend was observed in the Australian but not the Asian-Australian clades when modelled separately. In contrast to the evolutionary phenomena facilitating the development of reduced antibiotic susceptibility, adaptive evolution in the Australian clade appeared to be the sole contributor to enhanced CHX tolerance. The Asian-Australian clade is less tolerant to CHX, subsequently its introduction into the region, although increasing the prevalence of *qacA*, contributed minimally to the population level shift in phenotype (Table 2). In the Australian clade, the presence of a pSK1-like plasmid was significant associated with increase in CHX MIC and MBC (*p* < 0.0001, Table 2). However, an increase in the prevalence of these plasmids in the population alone could not account for the extent of intra-clade variation observed (Figure 4), and when the plasmid-harbouring population was modelled separately the trend towards enhanced tolerance remained (Figure S3). Comparison of the average MIC and MBC between the pSK1-like variants suggested the more recently emerged SVs (SV2 to SV6) may be associated with increased tolerance (Figure 5, Table S2). When the plasmid-harbouring population were grouped based on structural similarities, chromosomal integration was significantly associated with an increased CHX MIC (*p* = 0.0086, Table S2). Furthermore, a multi-CDS deletion in the plasmid backbone appeared to be associated with a further increase in MIC, although this was statistically non-significant (Table S2). It is unclear why these structural changes were only associated with a shift in MIC, with all groups having demonstrated highly consistent average MBC values (Figure 5), but this might reflect the specific mechanisms mediating enhanced CHX tolerance or the method utilised to estimate the MBC.

**Figure 4.**
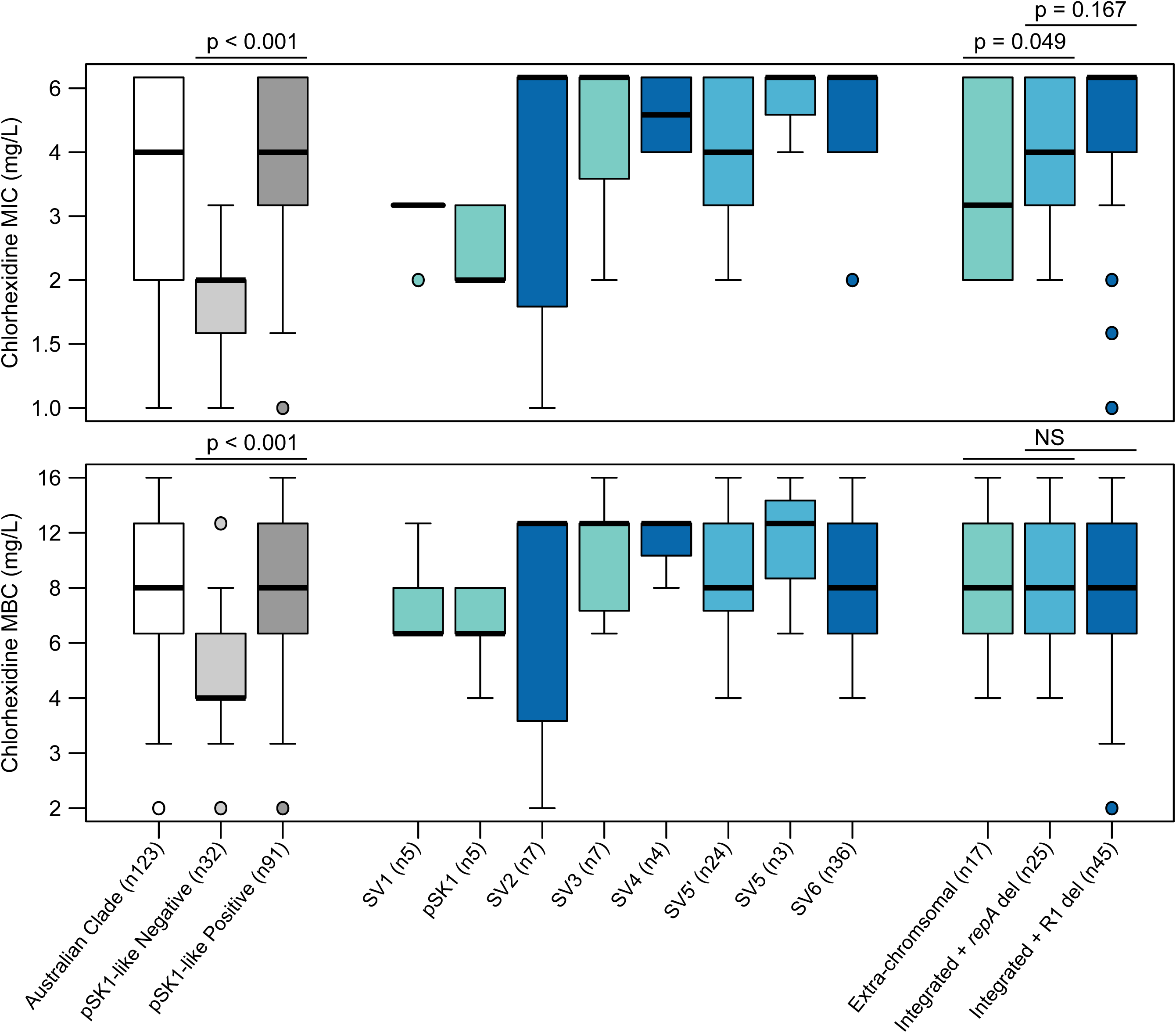
Phenotypic variation in chlorhexidine tolerance. Graphs illustrate the distribution of chlorhexidine MIC (top panel) and MBC (bottom panel) values in the Australian clade. Boxplot features represent the population median (central black line), upper and lower quartiles (box), and range (bars) excluding outliers (circles). Boxplots representing SVs are coloured to reflect a plasmid structural feature: extra-chromosomal plasmid (teal), and chromosomally integrated with either an internal *repA* deletion (light blue) or a multi-CDS region 1 deletion (dark blue).

**Figure 5.**
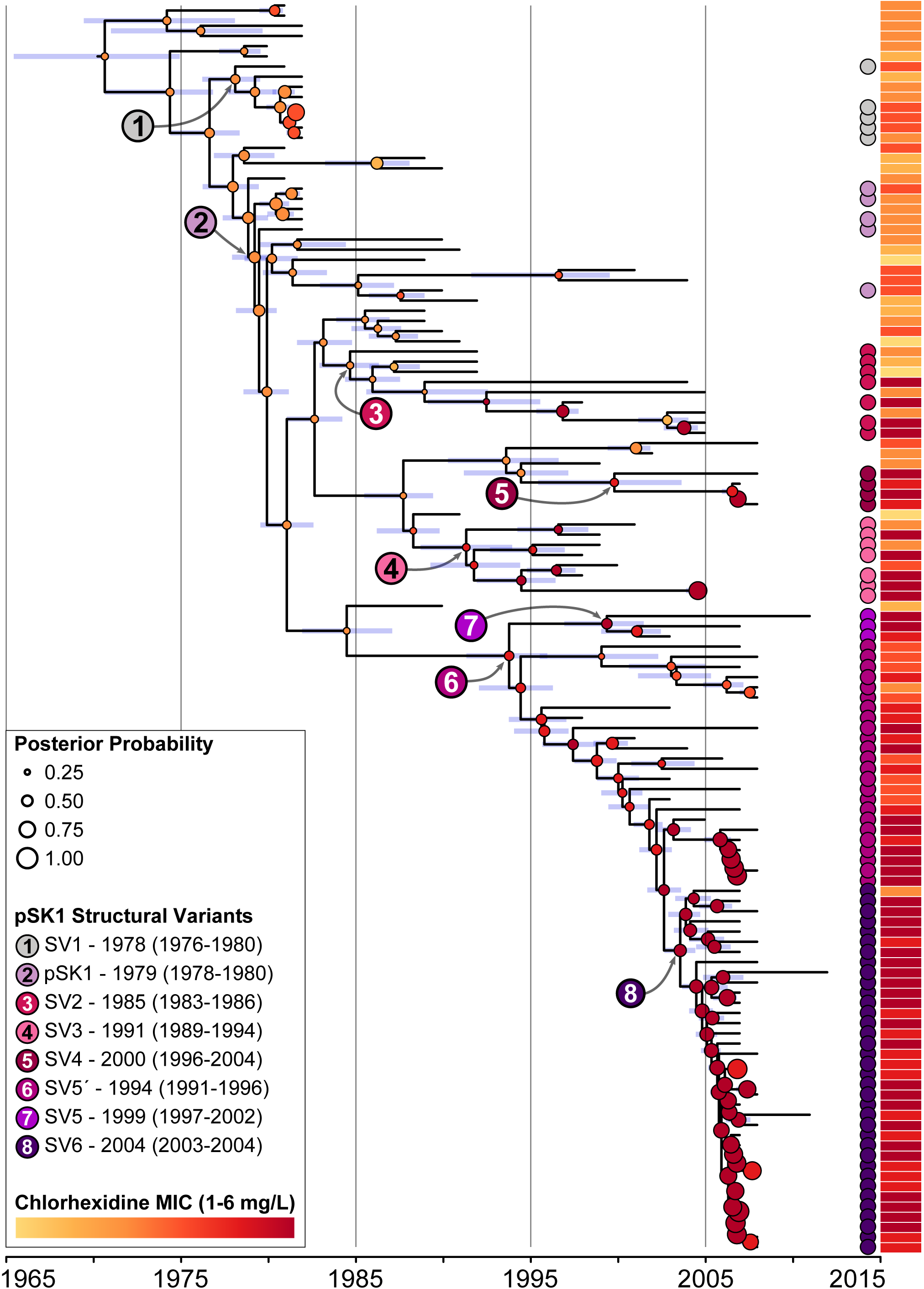
Bayesian phylogenetic model associating chlorhexidine tolerance with pSK1-like plasmid evolution. Illustrated is a maximum clade credibility tree inferred from the whole genome alignment of the Australian clade (n = 124). Isolates identified as harbouring a pSK1-like plasmid are indicated by a circle located adjacent to the tree and coloured based on the SV identified. The ancestral nodes in which each SV is estimated to have emerged are indicated by a number (refer to key), those coloured black represent an extra-chromosomal plasmid and those coloured white represent a genomic island. The estimated CHX MIC for all ancestral nodes is indicated by a circle, coloured based on the MIC value and sized according to the posterior probability for the estimate (refer to key). Blue bars represent the 95% highest posterior density interval for the node heights. The aligned heatmap illustrates the phenotypic MIC values attained for each isolate.

Collectively, these findings suggest that convergent evolution of pSK1 is associated with and likely contributing to the development of enhanced CHX tolerance in the Australian clade. Subsequently CHX use, which is extensive in the healthcare environment, being a fundamental component of infection control practices and hand hygiene initiatives in Australia and proven to be effective in reducing rates of invasive staphylococcal disease [57, 58], is a possible driver of pSK1 evolution.Although strongly suggested by these data, this hypothesis will need additional experimentation to confirm this association and demonstrate that exposure to CHX promotes pSK1-like plasmid maintenance through chromosomal integration.

### Independent Association of CHX Tolerance with pSK1-Like Plasmid Evolution

To examine the strength of the association between plasmid evolution and the development of enhanced CHX tolerance in the ST239 population, in addition to exploring other possible mechanisms that may be contributing to this phenotype, we used three separate statistical-genomic techniques. In contrast to the previous analyses, these models have focused on the CHX phenotype with no prior assumption that plasmid evolution is a contributing factor.

Common approaches to associating phenotype with genotype involve techniques such as Genome Wide Association Studies (GWAS) and Discriminant Analysis of Principal Components (DAPC), which look for the presence/absence of mutations and/or genes that are disproportionately correlated with a phenotype of interest. Both approaches were utilised in this work, neither identifying any significant genotypes (outside of the *qacA*-harbouring plasmids) that could be responsible for enhanced CHX tolerance (Supplementary Results).

We therefore performed a modified Bayesian phylogeographic analysis, modelling the CHX phenotype rather than geographic location of the ancestral genomes. The hypothesis was that if the development of CHX tolerance was a consequence of plasmid evolution then the ancestral nodes (ANs) from which the SVs were estimated to have emerged should correlate with those in which a shift in MIC is predicted (Figure 5). In this model, a shift in MIC was predicted at the expected ANs for the emergence of the newer SVs. In the case of SV3, SV4 and SV5’ the estimated ancestral MIC had increased to 4 mg/L, from either 2 or 3 mg/L in the preceding ANs. The emergence of SV5 and SV6 also correlated with a shift in MIC from 4 to 6 mg/L. In the older SVs, a shift in MIC was modelled to have occurred a few nodes following the emergence of the SV. This discrepancy is likely due to the presence of multiple isolates not harbouring a pSK1-like plasmid in close proximity to these ANs. In the case of SV1, a shift in MIC from 2 to 3 mg/L was observed in the AN for a sub-clade of isolates. Similarly, a shift in MIC from 2 to 6 mg/L was observed in the AN for a sub-clade of SV2 harbouring isolates. The overall strong correlation between the emergence of the different pSK1-like SVs and estimated shifts in phenotype (5/8 ANs for all SVs, or 7/8 ANs when the SV1 and SV2 sub-clades are considered), in combination with the absence of an identifiable alternative genotypic mechanisms, provides further support for the association between structural remodelling in the pSK1-like plasmid population and the development of enhanced CHX tolerance in ST239 MRSA.

## Conclusions

Plasmids and other MGEs play a central role in the successful evolution and adaptation of bacterial populations but are often overlooked because of the challenges with examining some plasmid DNA sequences using short-read data. Here, we have provided a comprehensive analysis of the evolution of the pSK1-like plasmid population that has co-evolved with the Australian ST239 MRSA lineage for multiple decades. Within the ST239 MRSA population circulating in Australia, the pSK1-like plasmid population is structurally diverse, with eight distinct variants identified having arisen largely through *IS*256*-* and IS*257*-mediated loss/gain of the composite transposons, chromosomal integration, deletion/exclusion of CDS, and inversions. When assessed by a temporal phylogenetic model, it appeared that the plasmid population had undergone convergent evolution, with the repeated emergence of chromosomally integrated SVs. In investigating potential drivers for plasmid evolution, it was identified that chromosomal integration was strongly associated with the development of enhanced CHX tolerance. While the mechanism mediating enhanced tolerance remains unclear, we speculate that it is linked to altered regulation of the *qacAR* efflux system, potentially resulting from the movement of IS elements and changes in plasmid configuration. These findings support the idea that the widespread and increasing use of CHX is possibly contributing to the evolution of the pSK1 family of plasmids. Although the levels of reduced susceptibility observed in this study remain well below in-use concentrations for this biocide, they do illustrate an evolutionary response in ST239 MRSA, one that may provide an adaptive advantage in healthcare institutions.

## Materials and Methods

### Bacterial Isolates

This study utilised a temporal (recovered between 1980 and 2012) and geographically diverse collection of 212 Australian ST239 *S. aureus.* Two isolates, including reference *S. aureus* JKD6008, were recovered in New Zealand [59]. All isolates represented cases of clinical infection. To establish global phylogenetic context for the collection, we supplemented this data with the WGS data for a further 319 international ST239 *S. aureus.* Relevant isolate information can be found in Supplementary Dataset.

### Whole Genome Sequencing & Sequence Data

The WGS data for 368 of the 531 isolates (73 of the 212 Australian isolates) had been previously published. All sequence data novel to this study has been made publicly available. Seven isolates were subjected to long-read sequencing to enable complete genome assembly. Information about the generation of novel sequence data, relevant WGS information and accession numbers can be found in the Supplementary Materials.

### Bioinformatic Analysis

The bioinformatic analyses performed for this study have been explained in detail in the Supplementary Methods and are briefly outlined here. Sequence data was mapped to reference *S. aureus* JKD6008 (GenBank accession CP002120, [60]) or reference plasmid pSK1 (NC_014369, [31]) using Snippy v3.2 (https://github.com/tseemann/snippy). Maximum likelihood phylogenetic trees were generated with IQ-TREE v1.6.1 [61], and maximum clade credibility trees were generated with BEAST v2.4.7 [62]. Trees were visualised in FigTree v1.4.3 (http://tree.bio.ed.ac.uk/software/figtree/) and figures were assembled in Inkscape v0.91 (https://inkscape.org/). Short read sequence data was *de novo* assembled using SPAdes v3.11.0 [63], and annotated with Prokka v1.12 [64]. Long-read sequence data was *de novo* assembled using the SMRT Analysis System v2.3.0.140936 (Pacific Biosciences), circularised and reorientated in Geneious v8.1.5 (Biomatters), and polished with Snippy v3.2. Plasmid structural comparisons were conducted using the Artemis Comparison Tool [65]. Ortholog clustering was performed using Roary [66].

### Antimicrobial Susceptibility Testing

Phenotypic susceptibility testing to trimethoprim and gentamicin was performed using E-tests (bioMérieux), and interpreted using CLSI guidelines (M100S, 26^th^ Ed). Susceptibility to CHX was performed using a modified broth microdilution method. CHX MICs were read after 24 hours incubation, and all wells were sub-cultured to assess viability. All isolates were tested in biological triplicate and the median values used for statistical analysis. Susceptibility testing procedures are explained in detail in the Supplementary Methods. All statistical analyses were performed in R v3.4.2 (http://www.R-project.org/), with significance determined as a p value ≤ 0.05.

## Acknowledgements

This work was funded by the National Health and Medical Research Council, Australia (Fellowships GNT1105905 to BPH; Project GNT457454 to NF and SOJ). SLB was supported by an Australian Government Research Training Program Scholarship, and a Victorian Fellowship provided by the Victorian State Government.

We thank Eloise Alison for her assistance in generating the phenotypic susceptibility data.

## Supplementary Materials

**Supplementary Dataset.** This file summarises the relevant isolates demographics and WGS information (including sequence data accession numbers) for all isolates included in this study.

**Supplementary Methods.** This file contains additional information about materials and methods used in this study.

**Supplementary Tables.** This file contains additional tables which provide information about the distribution of gentamicin and chlorhexidine susceptibility in the ST239 MRSA population.

**Supplementary Results.** This file contains additional analyses and results relevant to this study.

**Supplementary Figure S1. Global population structure of ST239 *S. aureus.*** Illustrated is a maximum clade credibility tree inferred from the whole genome alignment of the 531 international ST239 isolates. Tips are coloured based on location (refer to keys). Nodes with > 95% posterior support are indicated by a red dot. The Australian and two Asian-Australian clades (major and minor) and estimates for the most recent common ancestor (MRCA) are indicated, displayed as “median year (95% highest posterior density range)”.

**Supplementary Figure S2. Phylogenetic model for the emergence of pSK1 plasmid variants.** Illustrated is a maximum clade credibility tree inferred from the whole genome alignment of the Australian clade (n = 124). Isolates identified as harbouring a pSK1-like plasmid are indicated by a circle located at the branch tip and coloured based on the SV identified (refer to key). The most recent common ancestor (MRCA) for each SV is indicated by the larger node circle, with black numbers indicating an extrachromosomal SV and white numbers a chromosomally integrated SV. Temporal estimates for these nodes have been provided, displayed as “SV – median year (95% highest posterior density range)”. The two isolates with pSK1 region 1 (R1) deletions in the SV5’ and SV5 clade are indicated by a star. Grey arrows illustrate the likely order of structural remodelling that has occurred during the evolution of the pSK1-like plasmid population. Branches with < 95% posterior support are coloured red. The 95% highest posterior density for node heights is represented by the blue bars.

**Supplementary Figure S3. Modelling temporal-association in phenotypic susceptibility data.** Graphs depict linear models developed to explore the potential association between gentamicin MIC and chlorhexidine MIC and MBC with the year in which isolates were recovered. The dotted line indicates the smoothed mean MIC and the bold line indicates the fitted linear model. Four populations were tested (from top to bottom): (i) All ST239 MRSA (n = 211), (ii) the Asian-Australian clade (n = 88), (iii) the Australian clade (n = 123), and (iv) the pSK1-like plasmid harbouring population (n = 96).

